# Protein model quality assessment using 3D oriented convolutional neural networks

**DOI:** 10.1101/432146

**Authors:** Guillaume Pagès, Benoit Charmettant, Sergei Grudinin

## Abstract

Protein model quality assessment (QA) is a crucial and yet open problem in structural bioinformatics. The current best methods for single-model QA typically combine results from different approaches, each based on different input features constructed by experts in the field. Then, the prediction model is trained using a machine-learning algorithm. Recently, with the development of convolutional neural networks (CNN), the training paradigm has changed. In computer vision, the expert-developed features have been significantly overpassed by automatically trained convolutional filters. This motivated us to apply a three-dimensional (3D) CNN to the problem of protein model QA.

We developed a novel method for single-model QA called Ornate. Ornate (Oriented Routed Neural network with Automatic Typing) is a residue-wise scoring function that takes as input 3D density maps. It predicts the local (residue-wise) and the global model quality through a deep 3D CNN. Specifically, Ornate aligns the input density map, corresponding to each residue and its neighborhood, with the backbone topology of this residue. This circumvents the problem of ambiguous orientations of the initial models. Also, Ornate includes automatic identification of atom types and dynamic routing of the data in the network. Established benchmarks (CASP 11 and CASP 12) demonstrate the state-of-the-art performance of our approach among single-model QA methods.

The method is available at https://team.inria.fr/nanod/software/Ornate/. It consists of a C++ executable that transforms molecular structures into volumetric density maps, and a Python code based on the TensorFlow framework for applying the Ornate model to these maps.

## 1. Introduction

Proteins are ubiquitous for virtually all biological processes. Identifying their role helps to understand and potentially control these processes. However, even though protein sequence determination is now a routine procedure, it is often very difficult to use this information to extract relevant functional knowledge about system under study. Indeed, the function of a protein relies on a combination of its chemical and mechanical properties, which are defined by its structure.

Identifying protein structure from its sequence is thus a very important, though a challenging task. Experimental structure identification is not possible in all of the cases, and is generally very tedious and expensive. Therefore, computational methods that try to predict protein structure from its sequence have emerged in the past. Most of these methods combine the sampling of protein conformations step with the model quality assessment (QA) step. The former generates protein conformations, while the latter scores these to select the ones that will be as close as possible to the native structure.

In this work we only address the second problem and propose a novel method for protein model QA. This problem is challenging as it is shown by the fact that the Critical Assessment of protein Structure Prediction (CASP) community experiment (1) has an entire category dedicated to this specific topic (2). Indeed, the folding of a protein to its native conformation is driven by thermodynamic laws. This process can be formally characterised by the changes in free energy, which includes both enthalpic and entropic contributions. The former is defined by the potential energy contributions, while the latter describes the shape of the potential energy landscape. A proper estimation of the free energy differences is a very difficult and computationally expensive task, as it generally assumes knowledge about the protein environment, which is rarely available, and includes high-dimensional sampling.

Many methods for protein folding QA have already been developed. The goal of these methods is to predict the folding quality of a protein structure receiving its three-dimensional (3D) model as input. Generally, QA methods can be split into several classes. The best performing methods are often the consensus-based ones. This means that they do not score one single model but a whole set of them by comparing the models to each other. This class of methods is represented by Pcons (3) or 3D-Jury (4), for example. These methods are among the best performers on various benchmarks, but they suffer from the fact that one model cannot be scored alone and its score depends on the quality of other models in the scoring set. The methods that do not use consensus are called single-model methods. Among the single model methods, one can distinguish simple methods such as VoroMQA (5) or RWplus (6), which rely on a single type of structural features (contact area or pairwise atomic distances, respectively). Composite methods such as SBROD (7) aggregate many types of heterogeneous structural features. Meta-methods such as QProb (8) or DeepQA (9) integrate results from different methods to obtain better results. The boundaries between these categories are not always clear as some methods like Proq3D (10) aggregate both structural features and Rosetta energy terms (11).

The advent of machine learning techniques together with the growing amount of known 3D protein structures have broaden our possibilities to construct novel model QA methods. Specifically, convolutional neural networks (CNN, also sometimes referred to as deep learning) have demonstrated outstanding capacities for learning hierarchical representations (12). Very recently, 3D CNN has been applied to prediction of protein binding sites and also their interactions with ligands (13–16) and also protein QA tasks (17). However, one of the major hurdles for the success of 3D CNNs in this topic has been the uncertainty of choosing the reference orientation for the structures in the training and the test sets. For example, to circumvent this problem, (17) had to significantly augment the training set with random orientation of the input structures in 3D.

This work reports on a significant improvement of the 3D CNN method applied to the protein QA task, mostly due to the found solution to the orientation problem of the input 3D data. To do so, first, we decompose the global QA scoring task into a set of residue-wise local scoring tasks. Second, each local scoring task is handled by a residue-wise CNN with the input data oriented according to the local backbone topology. Other improvements of our 3D CNN model include automatic identification of atom types and dynamic routing of the data in the network. The performance of our model in the model QA task surpasses other single-model methods that rely only on the structure of the model. It is also very close to the performance of composite meta-methods that also use sequence alignment and evolutionary information as additional features (18).

## 2. Method

### A. Residue-wise scoring

Our model, called Ornate, which stands for Oriented Routed Neural network with Automatic Typing, relies on predicting local quality measures for each residue in a protein, provided a density map of its neighbourhood. There are several advantages of such technique compared to scoring the whole protein at once. Firstly, one protein structure contains many residues that provide multiple 3D examples to be used in training. Since the predicted score is local, the network can specifically learn local favoured or undesirable 3D geometries. This would not be possible by predicting the global score. Secondly, convolutional neural networks traditionally use a fixed input size. However, there are orders of magnitudes between the sizes of the smallest and the biggest proteins. Thus, choosing a fixed input size for the whole protein implies either constructing an oversized network, which will be costly to train and to run, or to be limited by the size of the structure to score. We should also notice that scaling the input structure to the input size of the network is very undesirable in our case, since we expect our network to learn some features that are not scale-invariant (e.g. hydrogen bonds or secondary structure). Finally, Ornate naturally provides a score per protein residue. This can be valuable for certain applications, where local scores are required. To obtain the global score for the overall model QA task, one simply needs to combine all the predicted local scores. This can be done in multiple ways. The score given to a structure by Ornate is the mean of the scores given to each of its residues.

There are, generally, multiple options for ground-truth local quality measures (19). For our purposes we required a relatively fast residue-wise measure. Therefore, we considered CAD-score (20) and LDDT (21), and have chosen the former. This choice was motivated by the smoothness of CAD-score, and the absence of an arbitrary score threshold, even though the two scores are very correlated (19).

### B. Input

For the estimation of the residue-wise score, Ornate is trained on cubic volumetric maps of side 19.2 Å centred and oriented on the given residue. This way, each map represents a certain residue with its spatial neighbourhood. Figure 1 shows an example of the protein volume captured by such a map.

**Fig. 1.**
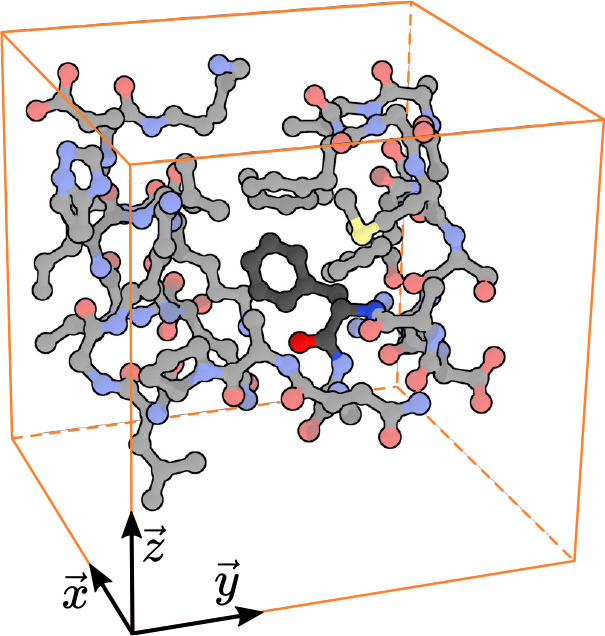
Example of the volumetric input corresponding to one protein residue (here, Phe58 from the 1yrf structure). The atoms of the considered residue are shown in dark colors and the atoms of his neighbourhood are shown in light colors. The orange box shows the boundaries of the considered neighbourhood. Only the atoms within this neighbourhood are shown.

#### Input orientation

Formally, the *n^th^* residue with its neighbourhood is represented by a cubic map positioned and oriented according the positions of its backbone atoms. The 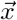 direction of the map coincides with the vector pointing from the carbon of the previous residue (*C*_*n*−1_) to the nitrogen of the current residue (*N*_*n*_). For the first residue, we define 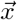 with the vector pointing from the alpha carbon 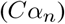 of the current residue to *C*_*n*_. The 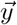 direction of the map is perpendicular to 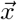 and is defined such that 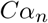 lies in the halfplane 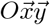 with *y* > 0 (see Fig. 2). Finally, 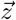 is defined as a vector product, 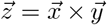. Once the three basis vectors are defined, we specify the origin of the map such that *N*_*n*_ is located at (6.1 Å, 6.6 Å, 9.6 Å) with respect to the map origin. This position has been chosen empirically such that all the atoms of all the residues among tested proteins fit in the map. This way, by definition, the position of *N*_*n*_ in the local frame is always the same, *C*_*n−*1_ is constrained along the 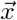 axis and 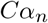 is constrained in the 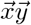 plane. In addition to these construction restraints, positional variance of other backbone atoms is also significantly reduced thanks to the fact that the values of the bond length and bond angles do not vary much. Also, a double bond between *N*_*n*_ and *C*_*n−*1_ forces the alpha carbon and the oxygen of the previous residue (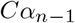 and *O*_*n−*1_) to lie in the same plane as *N*_*n*_, *C*_*n−*1_ and 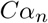. We specifically designed the local frame to keep constant the positions of as many atoms as possible. Indeed, we believe that having some invariant patterns allows CNN learning input structure better and faster, similarly to a situation with a human’s brain, which recognizes characters from a picture more easily if the picture is oriented correctly. By explicitly defining the origin and orientation of the input map with respect to the backbone atoms of the residue, we do not need the network to be rotationally invariant (22) and no data augmentation by rotating or translating the input is required (23).

**Fig. 2.**
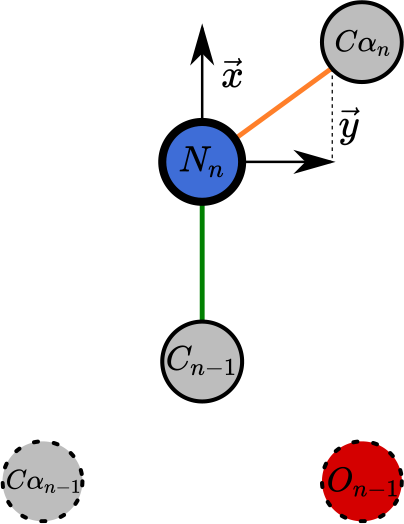
Illustration of the definition of the local frame for the *n*^th^ residue and the atoms positionally constrained by this definition. The local frame is defined by the position of *N*_*n*_ (blue), and the directions of *C*_*n*−1_ and 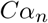 with respect to *N*_*n*_ (respectively shown in green and orange). *N*_*n*_ atom is shown with a bold outline, and its position is fixed in the local frame. Thin outlines correspond to the atoms whose positions are partially constrained in space. Dashed outlines correspond to the atoms whose positions are unconstrained, but in practice do not vary much from the mean values.

#### Input values

The input maps are constructed from the atomic representation of the position of the current residue and its neighbouring atoms. The atomic representation of the structure is first transformed to a density function, then projected on a grid to obtain the map input for our CNN. The density function associates each point in space with a vector of 167 dimensions. These 167 dimensions correspond to the 167 different atom types that can be found in amino acids (without the hydrogens). A list of these atom types is given in Supplementary Information. More formally, let 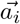 be the position of the *i*^*th*^ atom of the structure, *σ* be the width of the Gaussian kernel (we use *σ* = 1Å) and *t*_*i*_ be the 167-dimensional unit vector whose only non-zero component corresponds to the type of the *i*^*th*^ atom. The density function ***d*** at a point 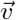 then reads as

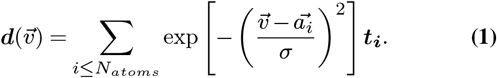

To project the density on a map, we split the map in 24×24×24 voxels of side 0.8 Å, and assign to each voxel the value of the density function at its centre. Figure 3 shows an example of the projection made with three atoms with different atom types, represented by red, green and blue colors. To reduce memory footprint, we store each component of a voxel value in one byte of data as a fixed-point number with a scaling value of 1/255. Thus, the map for one residue requires 24 * 24 * 24 * 167 = 2.3 MB of memory storage. As a consequence, a density smaller than 1/255 will be regarded as zero. This naturally truncates to zero values of the Gaussian kernel with arguments larger than 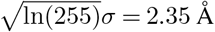. It is thus possible not to consider atoms that are more distant than 2.35 Å from the map, and, with an appropriate neighbour search algorithm, to keep a linear complexity of the mapping algorithm with respect to the number of residues in the protein.

**Fig. 3.**
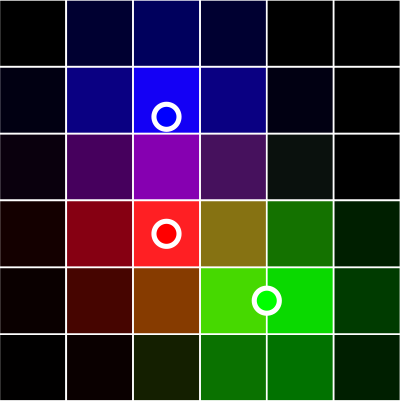
Example of three atoms projected on a map with the presented method. For clarity, we only show 6 × 6 voxels of a 2D slice of the map. Three atoms of different atom types are represented by three circles of different colors. The figure is scaled so that the side of one voxel is 0.8 Å, and the inter-atomic distance is 1.4 Å, which is a typical bond length for heavy atoms in proteins.

### C. Network topology

The CNN architecture used in this work is inspired by CNNs from computer vision. Figure 4 summarizes the network’s topology. A typical CNN design begins with convolutional layers that deal with high dimensions of the input spatially-structured data, followed by fully connected layers after the dimensionality of the data has been reduced. In our design, the three convolutional layers learn high-level features while progressively coarsen them and reducing the data’s dimensionality. Then, a set of fully connected layers combines these features and outputs a prediction. As the activation function, we used ELU (24), which has been proven to speed up learning in deep neural networks and lead to higher classification accuracies. We also include batch normalization layers (25) that accelerate the training and improve the accuracy of the network’s predictions. Finally, we added two additional layers, a “retyper” and a “router” designed for specific purposes, which are explained below.

**Fig. 4.**
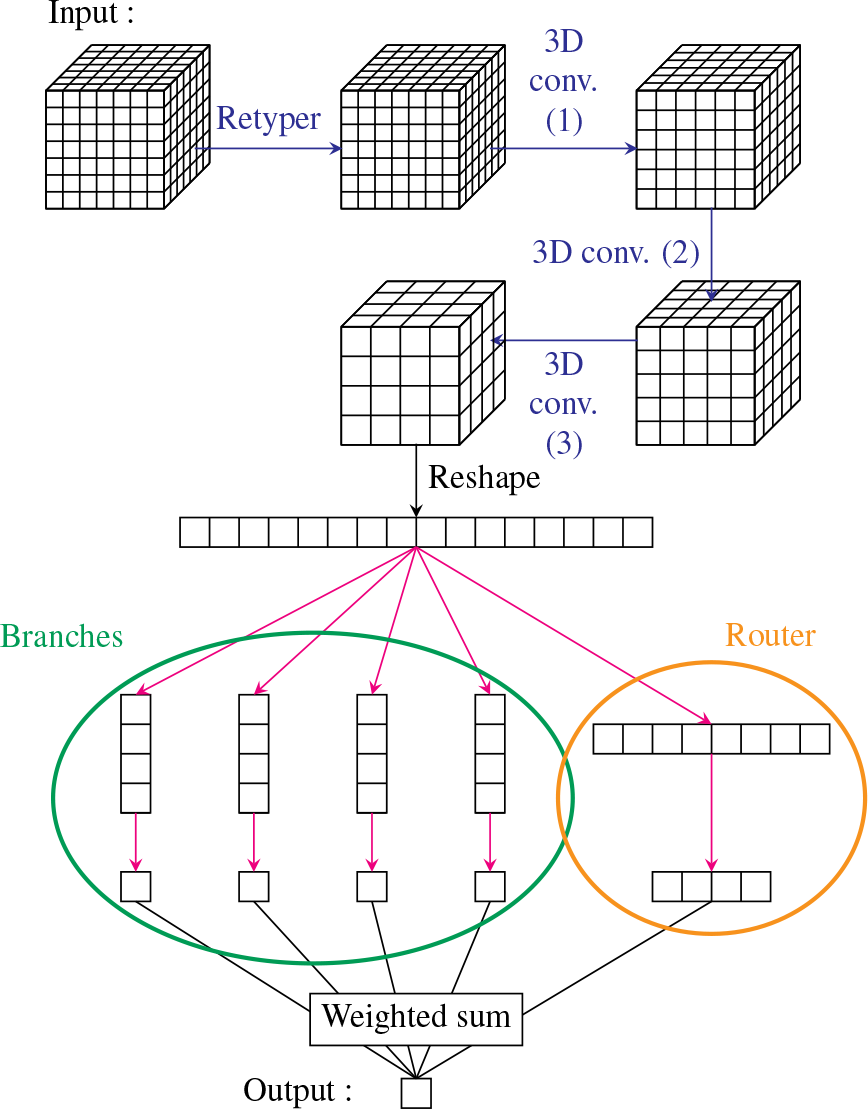
Summary of the network topology. The blue arrows represent 3D convolutional layers and the red arrows represent fully connected layers. A more complete description is given in Supplementary Information.

#### The retyper layer

A very original and uncommon pattern of the current network is the presence of the first layer that we called “retyper”. This layer, which is technically a convolutional layer of size 1 × 1 × 1 with 167 input channels and 15 output channels, projects each of the 167 atom types that exist in proteins to a feature space of dimension 15. By doing so, we reduce the dimensionality of the input by a factor of 11, and switch from a sparse data representation to a dense one. Indeed, a voxel with *n* nonzero components implies *n* atoms of different types located at a distance smaller than 2.35 Å from the voxel center. In practice, there can be at most a dozen of nonzero components in one voxel.

#### Router layer

After the convolutional layers, we apply a data routing layer. Our initial idea was to explicitly allow the combination of the features to be different depending on the residue type provided in input. We separately trained 20 different routes as a second part of our network, which were specific to each type of amino acids. However, to do so, we needed 20 times more training steps because only one route was trained at a time. In practice, some routes should require even more training steps, since the amino acid distribution is not even in proteins. Altogether, the gain from having a different model for each route was not worth the additional training.

As a second attempt, we later changed our network architecture to let the network learn the data routing as proposed in (26). The idea here was to have a network called “router” that predicts which route should be trusted to score these particular data. In this implementation, the data outputed by the convolutional layers are sent to every route and the final score is an average of the different outputs, weighted by the router predictions. The advantage compared to the previous technique is that the router can learn more relevant criteria than just the residue type to choose which route to select.

### D. Training loss function

We chose to train Ornate to approximate the value of CAD-score (20) of each residue. As a result, we set the training loss function for a residue *r*_*i*_ scored *s*(*r*_*i*_) by our network as:

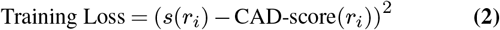

The training loss is thus simply the squared difference between the prediction and the ground truth for each residue.

### E. Training phase

As the training set, we used the server submissions for CASP 7, 8, 9, 10 stage 2 experiments. We also removed a few structures whose CAD-score were equal to zero, or whose backbone was incomplete. We trained Ornate with 100,000 optimization steps using a stochastic gradient descent method (please see Supplementary Information for more details). Each step optimizes the network on 10 consecutive residues from an input structure. When a structure is running out of residue, a new one is randomly selected. Thus, we used a total of 1M residues for optimizing the network. This represent less than 10,000 structures, while stages 2 of CASP 7, 8, 9, 10 contain each about 10,000 models. Each structure is thus used at most once, meaning that the values of the training loss function were always computed for new structures.

Figure 5 shows the training loss during each step of the training. Since this loss is very fluctuant, we also plot a smoothed version of the loss defined at each training step *n* as:

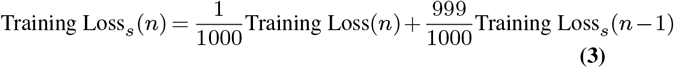

This smoothed version seems to reach a plateau after about 80,000 steps so we decided to stop the training at this point. The overall decrease of loss cannot be due to over-fitting, since each step was trained on new data.

**Fig. 5.**
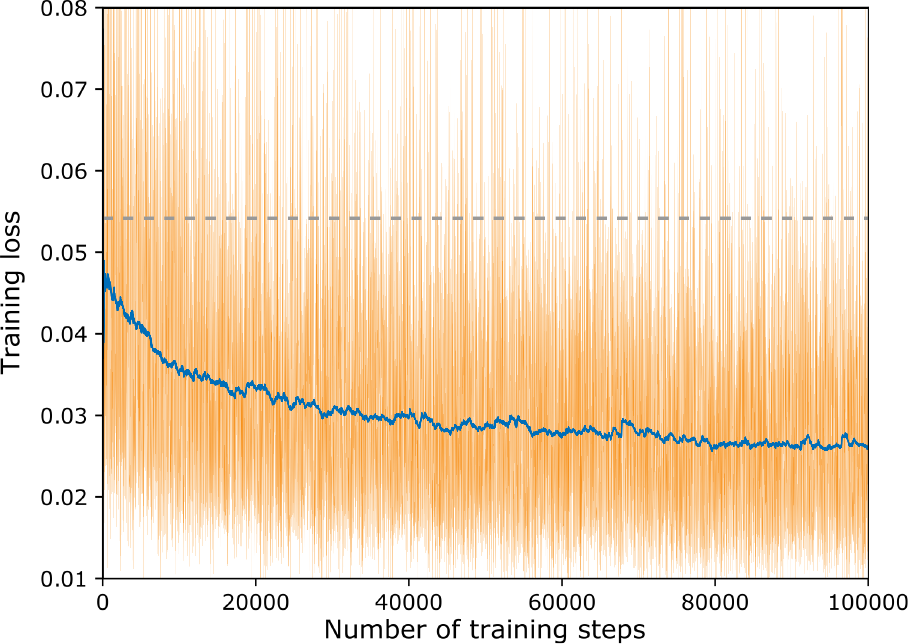
Variation of the loss during 100,000 training steps. The training loss is shown with the thin orange line and the smoothed loss with the thick blue line. The grey dashed line shows the variance of CAD-score on the training set. It equals to the expected value of the training loss for a scoring scheme that always returns the average CAD-score of the training set.

## 3. Results and discussion

### A. Comparison with the state-of-the-art

We compared the results of our scheme with several other state-of-the-art QA methods. To do so, we used the same benchmark as (7) and (8). For a rigorous comparison, we trained Ornate with the data from CASP 7 - 10 server submissions, and blindly scored protein models from CASP 11 and CASP 12 server submissions, stage 1 and 2. Formally, for each target of CASP 11, and CASP 12, we have a model set *M* = {*P*_1_,…, *P*_*n*_ }and a native structure *P*_0_. We computed estimators of the performance of QA methods *Q* with respect to the ground-truth measure *G*, where *G*(*P*) measures the similarity between the model *P* and native structure *P*_0_. The prediction loss (P. L.) is defined as:

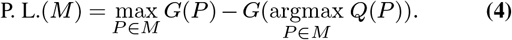

Let us define the average of a function *F* over a set *M* = {*P*_1_,…, *P*_*n*_} as

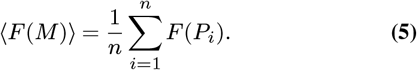

Pearson’s *r* is defined as

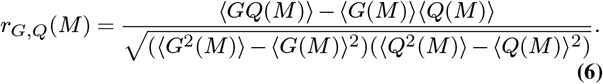

Spearman’s *ρ* measures the correlation between the rankings given by two functions

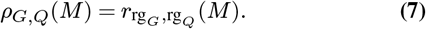

Kendall’s *τ* is defined by

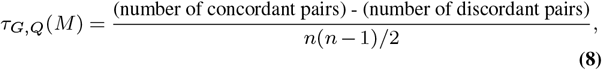

where a pair {*P*_*i*_, *P*_*j*_} is concordant if (*G*(*p*_*i*_) − *G*(*p*_*j*_)(*Q*(*p*_*i*_) − *Q*(*p*_*i*_)) > 0 and discordant if (*G*(*p*_*i*_) − *G*(*p*_*j*_)(*Q*(*p*_*i*_) − *Q*(*p*_*i*_)) < 0.

The average losses, Pearson’s *r*, Spearman’s *ρ* and Kendall’s *τ* are an average of these indicators computed for each target. They estimate how well the scoring function can compare structures with the same sequence, and how well it picks the best among them. In addition, we computed global Pearson’s *r*, Spearman’s *ρ* and Kendall’s *τ* on the union of all decoy sets. These estimate the capability of a method to compare quality of structures with different sequences, and thus to predict if a model is far or close to the native structure. We compared our method only with single-model methods (we excluded consensus-based methods), which are listed below. SBROD (7) is a coarse-grain knowledge-based method trained using four types of structural features, residueresidue pairwise features, backbone atom-atom pair wise features, hydrogen bonding features and solvent-solvate features. We used the version from https://gitlab.inria.fr/grudinin/sbrod trained on CASP 5-10 server predictions. VoroMQA (5) is a statistical potential trained on inter-atomic contact areas. We used the version included in the package voronota version 1.18.1877 from https://bitbucket.org/kliment/voronota/downloads/. RWplus (6) is a classical gold-standard statistical potential that uses a pairwise-dependent atomic potential, with a side chain orientation-dependent energy term. We used the binary provided at https://zhanglab.ccmb.med.umich.edu/RW/. 3DCNN (17) is another method that uses convolutional networks trained on protein density maps. We were not able to run this method on our side so we used the scores provided in http://proteinfoldingproject.com/static/datasets/models.tar.gz. Only the scores of CASP 11 are available in this archive. Proq3D (10) is a method that combines many heterogeneous expert-defined features using a deep-learning algorithm. It contains different back-end models to fit different geometrical scores. To fairly compare Proq3D with our method, we used the version of Proq3D trained on CAD-score with rotameric optimization enabled. We should also mention that Proq3D does not only rely on protein structure, as it uses a sequence database to extract the relevant evolutionary information (18). We used the method available at https://bitbucket.org/ElofssonLab/proq3 with a version of UniRef sequence database (27) from 2013 that was already available when CASP 11 challenge started.

**Table 1.** Performance of different methods for model quality assessment on CASP 11 and CASP 12 benchmarks. The ground-truth measure is CAD-score. The sign * indicates that the scoring function has been specifically trained to fit this measure. The three best performing methods are highlighted in orange with increasing saturation. Av. and Gl. stand for average and global. P.L., *r*, *ρ* and *τ* stand for prediction loss, Pearson’s *r*, Spearman's *ρ* and Kendall’s *τ*, respectively.

| Set | St. | Model | Av. P.L. | Av. $r$ | Av. $\rho$ | Av. $\tau$ | Gl. $r$ | Gl. $\rho$ | Gl. $\tau$ |
| --- | --- | --- | --- | --- | --- | --- | --- | --- | --- |
| CASP 11 | 1 | Ornate* | 0.030 | 0.700 | 0.641 | 0.501 | 0.788 | 0.771 | 0.577 |
|  |  | SBROD | 0.042 | 0.675 | 0.601 | 0.460 | 0.589 | 0.566 | 0.393 |
|  |  | VoroMQA | 0.036 | 0.687 | 0.600 | 0.460 | 0.728 | 0.725 | 0.532 |
|  |  | RWplus | 0.056 | 0.630 | 0.557 | 0.418 | 0.168 | 0.157 | 0.101 |
|  |  | 3dCNN | 0.059 | 0.408 | 0.409 | 0.310 | 0.487 | 0.521 | 0.359 |
|  |  | Proq3D* | 0.021 | 0.816 | 0.762 | 0.614 | 0.900 | 0.894 | 0.716 |
|  | 2 | Ornate* | 0.024 | 0.721 | 0.679 | 0.511 | 0.786 | 0.803 | 0.605 |
|  |  | SBROD | 0.043 | 0.578 | 0.550 | 0.392 | 0.548 | 0.560 | 0.385 |
|  |  | VoroMQA | 0.044 | 0.655 | 0.626 | 0.463 | 0.672 | 0.707 | 0.529 |
|  |  | RWplus | 0.053 | 0.418 | 0.419 | 0.300 | 0.108 | 0.097 | 0.063 |
|  |  | 3dCNN | 0.061 | 0.467 | 0.469 | 0.334 | 0.593 | 0.618 | 0.435 |
|  |  | Proq3D* | 0.024 | 0.714 | 0.687 | 0.511 | 0.885 | 0.891 | 0.710 |
| CASP 12 | 1 | Ornate* | 0.035 | 0.756 | 0.702 | 0.554 | 0.728 | 0.687 | 0.507 |
|  |  | SBROD | 0.029 | 0.663 | 0.546 | 0.414 | 0.380 | 0.291 | 0.205 |
|  |  | VoroMQA | 0.030 | 0.665 | 0.558 | 0.422 | 0.570 | 0.569 | 0.397 |
|  |  | RWplus | 0.052 | 0.567 | 0.506 | 0.377 | 0.010 | -0.009 | -0.005 |
|  |  | Proq3D* | 0.023 | 0.807 | 0.753 | 0.595 | 0.832 | 0.799 | 0.608 |
|  | 2 | Ornate* | 0.028 | 0.781 | 0.747 | 0.574 | 0.808 | 0.786 | 0.597 |
|  |  | SBROD | 0.049 | 0.686 | 0.630 | 0.470 | 0.507 | 0.491 | 0.342 |
|  |  | VoroMQA | 0.048 | 0.731 | 0.694 | 0.528 | 0.638 | 0.661 | 0.499 |
|  |  | RWplus | 0.063 | 0.642 | 0.597 | 0.431 | 0.050 | 0.048 | 0.037 |
|  |  | Proq3D* | 0.026 | 0.804 | 0.770 | 0.596 | 0.887 | 0.901 | 0.720 |

First, we evaluated the QA methods by comparing them with CAD-score as a ground-truth score. Table 1 lists the results. We can see that our method is almost always ranked second after Proq3D. We should note that historically, the protein structure prediction community was biased towards using GDT-TS (28) as a quality measure. Therefore, many of the machine learning-based methods (including those from our performance benchmark) have been specifically trained to approximate GDT-TS. Therefore, for a fair comparison, we ran an additional test. Here, we computed the loss, Pearson correlation and Kendall’s *τ* using GDT-TS as the groundtruth score. Table 2 lists the performance results. We can see that Proq3D is again the best performing method (even though it was trained on CAD-score). Here, our method performs, as expected, less impressive compared to the previous test (comparison with CAD-score). In particular, the average correlations are not as high as for the VoroMQA or SBROD methods, which were specifically trained to match GDT-TS. However, it is the second best method for picking the best structure in 3 out of 4 datasets, and it is also among the best for the global correlations, meaning that our method is still useful to assess the absolute quality of models. It is also very interesting to see the performance comparison between Ornate and 3dCNN. Indeed, 3dCNN is the most similar method to our work, and it was our starting point. Thus, performing better than 3dCNN demonstrates the progress we made with the choice of the network architecture. Our method performs better on every single indicator, in both CAD-score and GDTTS tests, even though 3dCNN has been specifically trained to match GDT-TS.

**Table 2.**
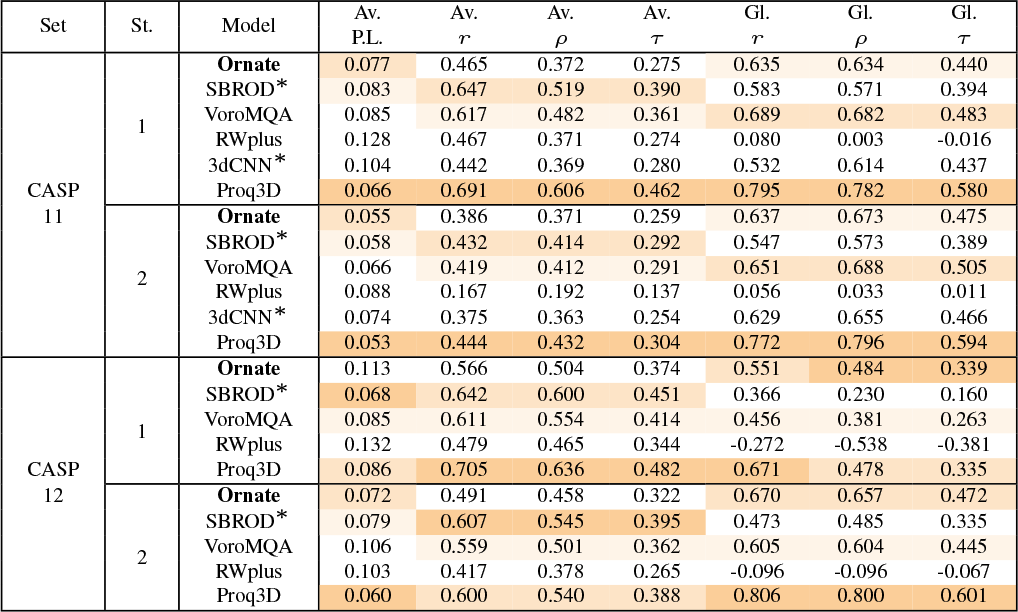
Performance of different methods for model quality assessment on CASP 11 and CASP 12 benchmarks. The ground-truth measure is GDT-TS. The sign * indicates that the scoring function has been specifically trained to fit this measure. The three best performing methods are highlighted in orange with increasing saturation. Av. and Gl. stand for average and global. P.L., *r*, *ρ* and *τ* stand for prediction loss, Pearson’s *r*, Spearman's *ρ* and Kendall’s *τ*, respectively.

### B. Local scores

A particularity of Ornate is to also compute local scores for each model residue. This helps to predict which part of the model structure is poorly folded or should be refined. Figure 6 shows a few examples of CASP server predictions scored with Ornate (for targets T0854, T0367, and T0768) with some correctly modeled parts and poorly modeled ones. As a reference, we also show the same models colored according to the ground-truth CAD-scores, and the reference crystallographic structures. This figure demonstrates a very good correlation between what Ornate predicts as poorly folded residues and what are actually the poorly folded residues.

### C. Computational details

We implemented the Ornate method using a combination of C++ and Python programming languages. The part for the generation of input volumetric maps was written in C++. The NN’s training part uses Python with the TensorFlow framework (29). The computational time thus can also be decomposed into two parts that take approximately the same time. Creating a 3D map from a residue structure and its neighborhood takes about 30 ms (measured with an I7 CPU), and running the network for one map takes about 20 ms (measured on a GeForce GTX 680 GPU). Please also note that the latter time was measured for TensorFlow with GPU support, and it may be up to 100 times slower without it. Overall, the complexity of scoring one protein model grows linearly with the number of residues in the model structure. For example, scoring a mid-size protein structure with about 200 residues takes about 1s.

## 4. Conclusion

This work presents Ornate, one of the first 3D CNN methods for the protein model QA problem. Ornate demonstrates a significant improvement over the previous 3D CNN attempts. This improvement was made possible thanks to several dedicated network topology designs that we introduced. These include residue-wise scoring, orientation of the input maps according to the backbone atoms, the retyper layer and data routing.

**Fig. 6.**
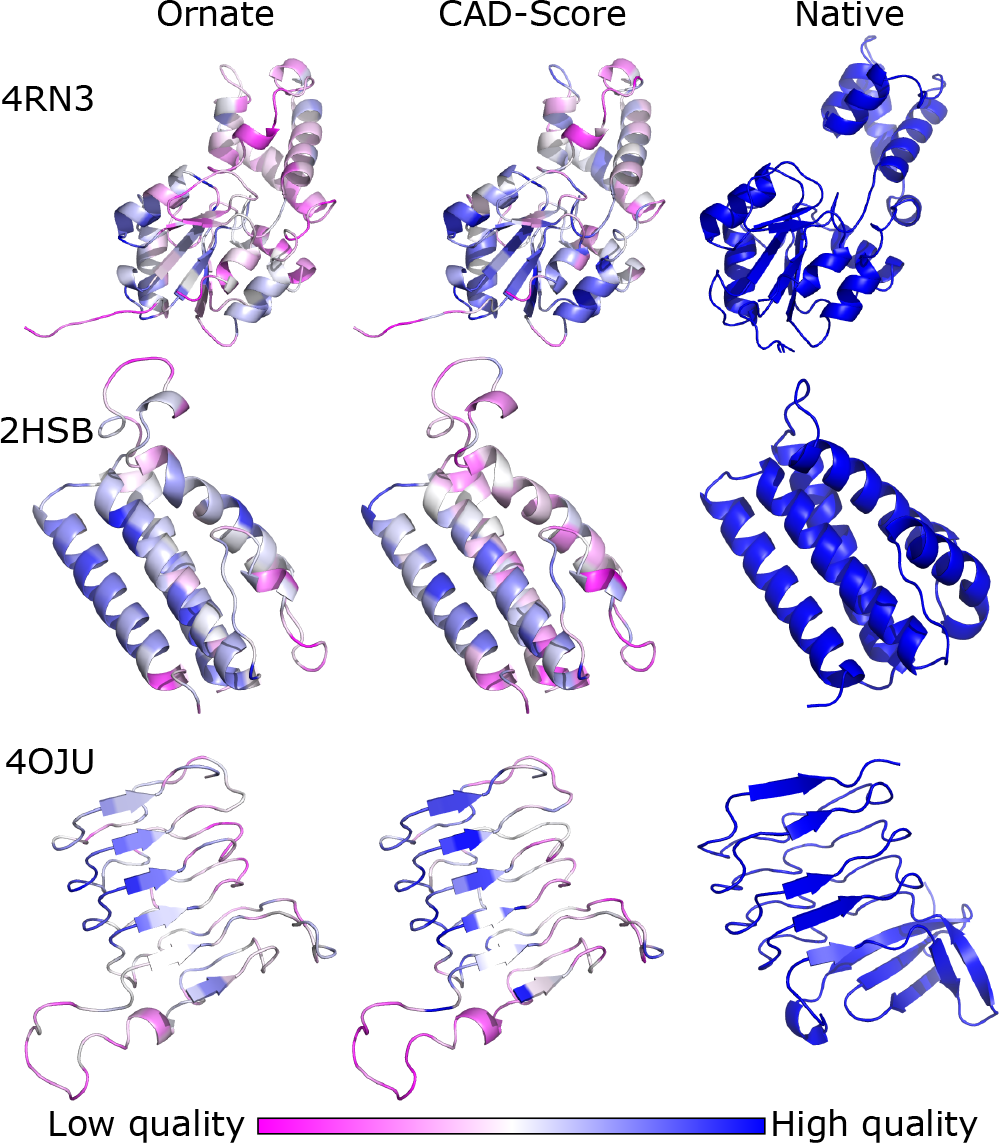
Examples of local quality measures for three CASP server prediction targets T0854 (top, pdb code 4rn3), T0367 (middle, pdb code 2hsb), and T0768 (bottom, pdb code 4oju). The left column shows models colored according to the Ornate score. The center column shows models colored according to the ground-truth CAD-score. The right column shows reference crystallographic structures. The three CASP models have respective GDT-TS measures of 0.657 (top), 0.634 (middle) and 0.469 (bottom), and CAD-score measures of 0.518 (top), 0.463 (middle) and 0.456 (bottom).

Ornate is competitive to most state-of-the-art single-model protein model QA methods. For example, when compared to the best of these, also trained to match CAD-score, on stage 2 of CASP 11 (which seems to be the hardest dataset to score according to our measures), the average correlations and prediction losses of Ornate are at the same level as the ones from Proq3D. However, Proq3D, compared to Ornate, accesses additional sequence data. Even though Ornate does not reach the accuracy of the best meta-methods on all the indicators, its reasonable scoring time (about 1 second for mid-size proteins) makes it a good candidate to be integrated in such metamodels. In addition, Ornate produces a smooth score with respect to atom positions in the model, as it has derivatives of all orders. Thus, its gradient can be easily computed. This property can be used for subsequent model refinement.

## 5. Acknowledgements

First, the authors thank Georgy Derevyanko from Concordia University, Montréal, who designed a preliminary version of a 3D CNN, which motivated us in constructing Ornate. The authors thank Kliment Olechnovic? from Vilnius University for his support on Voronota usage, and Arne Elofsson and Karolis Uziela from Stockholm University for their support on Proq3D usage. Finally, the authors thank Stephane Redon from Inria, Grenoble, for his permanent interest and motivated discussions in deep learning techniques, and also for his support on data visualization.

## Bibliography

1. John Moult, Krzysztof Fidelis, Andriy Kryshtafovych, Torsten Schwede, and Anna Tramontano. Critical assessment of methods of protein structure prediction (CASP)—Round XII. Proteins: Structure, Function, and Bioinformatics, 86:7–15, 2018.

2. Domenico Cozzetto, Andriy Kryshtafovych, Michele Ceriani, and Anna Tramontano. Assessment of predictions in the model quality assessment category. Proteins: Structure, Function, and Bioinformatics, 69(S8):175–183, 2007.

3. Jesper Lundström, Leszek Rychlewski, Janusz Bujnicki, and Arne Elofsson. Pcons: A neural-network–based consensus predictor that improves fold recognition. Protein Science, 10(11):2354–2362, 2001.

4. Krzysztof Ginalski, Arne Elofsson, Daniel Fischer, and Leszek Rychlewski. 3D-Jury: a simple approach to improve protein structure predictions. Bioinformatics, 19(8):1015–1018, 2003.

5. Kliment Olechnovič and Česlovas Venclovas. Voromqa: Assessment of protein structure quality using interatomic contact areas. Proteins: Structure, Function, and Bioinformatics, 85(6):1131–1145, 2017.

6. Jian Zhang and Yang Zhang. A novel side-chain orientation dependent potential derived from random-walk reference state for protein fold selection and structure prediction. PloS one, 5(10):e15386, 2010.

7. Mikhail Karasikov, Guillaume Pagès, and Sergei Grudinin. Smooth orientation-dependent scoring function for coarse-grained protein quality assessment. Unpublished, 2018.

8. Renzhi Cao and Jianlin Cheng. Protein single-model quality assessment by feature-based probability density functions. Scientific reports, 6:23990, 2016.

9. Renzhi Cao, Debswapna Bhattacharya, Jie Hou, and Jianlin Cheng. DeepQA: improving the estimation of single protein model quality with deep belief networks. BMC Bioinformatics, 17(1):495, 2016.

10. Karolis Uziela, David Menéndez Hurtado, Nanjiang Shu, Björn Wallner, and Arne Elofsson. ProQ3D: improved model quality assessments using deep learning. Bioinformatics, 33(10): 1578–1580, 2017.

11. Andrew Leaver-Fay, Michael Tyka, Steven M Lewis, Oliver F Lange, James Thompson, Ron Jacak, Kristian W Kaufman, P Douglas Renfrew, Colin A Smith, Will Sheffler, et al. Rosetta 3: an object-oriented software suite for the simulation and design of macromolecules. In Methods in enzymology, volume 487, pages 545–574. Elsevier, 2011.

12. Honglak Lee, Roger Grosse, Rajesh Ranganath, and Andrew Y Ng. Convolutional deep belief networks for scalable unsupervised learning of hierarchical representations. In Proceedings of the 26th annual international conference on machine learning, pages 609–616. ACM, 2009.

13. Izhar Wallach, Michael Dzamba, and Abraham Heifets. AtomNet: A deep convolutional neural network for bioactivity prediction in structure-based drug discovery. arXiv preprint arXiv:1510.02855, 2015.

14. Matthew Ragoza, Joshua Hochuli, Elisa Idrobo, Jocelyn Sunseri, and David Ryan Koes. Protein-ligand scoring with convolutional neural networks. Journal of chemical information and modeling, 57(4):942–957, 2017.

15. J Jiménez, S Doerr, G Martínez-Rosell, AS Rose, and G De Fabritiis. DeepSite: proteinbinding site predictor using 3D-convolutional neural networks. Bioinformatics, 33(19):3036–3042, 2017.

16. Raphael JL Townshend, Rishi Bedi, and Ron O Dror. Generalizable protein interface prediction with end-to-end learning. arXiv preprint arXiv:1807.01297, 2018.

17. Georgy Derevyanko, Sergei Grudinin, Yoshua Bengio, and Guillaume Lamoureux. Deep convolutional networks for quality assessment of protein folds. Bioinformatics, page bty494, 2018. doi: 10.1093/bioinformatics/bty494.

18. Arjun Ray, Erik Lindahl, and Björn Wallner. Improved model quality assessment using ProQ2. BMC bioinformatics, 13(1):224, 2012.

19. Kliment Olechnovicč, Bohdan Monastyrskyy, Andriy Kryshtafovych, Česlovas Venclovas, and Alfonso Valencia. Comparative analysis of methods for evaluation of protein models against native structures. Bioinformatics, 2018.

20. Kliment Olechnovicč, Eleonora Kulberkyteė, and Česlovas Venclovas. CAD-score: A new contact area difference-based function for evaluation of protein structural models. Proteins: Structure, Function, and Bioinformatics, 81(1):149–162, 2013.

21. Valerio Mariani, Marco Biasini, Alessandro Barbato, and Torsten Schwede. lDDT: a local superposition-free score for comparing protein structures and models using distance difference tests. Bioinformatics, 29(21):2722–2728, 2013.

22. Daniel E Worrall, Stephan J Garbin, Daniyar Turmukhambetov, and Gabriel J Brostow. Harmonic networks: Deep translation and rotation equivariance. In Proc. IEEE Conf. on Computer Vision and Pattern Recognition (CVPR), volume 2, 2017.

23. David A Van Dyk and Xiao-Li Meng. The art of data augmentation. Journal of Computational and Graphical Statistics, 10(1):1–50, 2001.

24. Djork-Arné Clevert, Thomas Unterthiner, and Sepp Hochreiter. Fast and accurate deep network learning by exponential linear units (elus). arXiv preprint arXiv:1511.07289, 2015.

25. Sergey Ioffe and Christian Szegedy. Batch normalization: Accelerating deep network training by reducing internal covariate shift. arXiv preprint arXiv:1502.03167, 2015.

26. Yani Ioannou, Duncan Robertson, Darko Zikic, Peter Kontschieder, Jamie Shotton, Matthew Brown, and Antonio Criminisi. Decision forests, convolutional networks and the models inbetween. arXiv preprint arXiv:1603.01250, 2016.

27. Baris E Suzek, Hongzhan Huang, Peter McGarvey, Raja Mazumder, and Cathy H Wu. Uniref: comprehensive and non-redundant UniProt reference clusters. Bioinformatics, 23 (10):1282–1288, 2007.

28. Yang Zhang and Jeffrey Skolnick. Scoring function for automated assessment of protein structure template quality. Proteins: Structure, Function, and Bioinformatics, 57(4):702–710, 2004.

29. Martín Abadi, Paul Barham, Jianmin Chen, Zhifeng Chen, Andy Davis, Jeffrey Dean, Matthieu Devin, Sanjay Ghemawat, Geoffrey Irving, Michael Isard, et al. Tensorflow: asystem for large-scale machine learning. In Proceedings of the 12th USENIX conference on Operating Systems Design and Implementation, pages 265–283. USENIX Association, 2016.

